# Revealing missing isoforms encoded in the human genome by integrating genomics, transcriptomics and proteomics data

**DOI:** 10.1101/012112

**Authors:** Zhiqiang Hu, Hamish S. Scott, Guangrong Qin, Guangyong Zheng, Xixia Chu, Lu Xie, David L. Adelson, Bergithe E Oftedal, Parvathy Venugopal, Milena Babic, Christopher N Hahn, Bing Zhang, Xiaojing Wang, Nan Li, Chaochun Wei

**Affiliations:** School of Life Sciences and Biotechnology, Shanghai Jiao Tong University, 800 Dongchuan Road, Shanghai 200240, China; Shanghai Center for Bioinformation Technology, 1278 Keyuan Road, Pudong District, Shanghai 201203, China; Department of Genetics and Molecular Pathology, Centre for Cancer Biology, SA Pathology, Adelaide, South Australia, Frome Road, Adelaide, SA 5000 Australia; School of Molecular and Biomedical Science, University of Adelaide SA 5000, Australia; School of Medicine, University of Adelaide, North Terrace, Adelaide, SA 5000, Australia; School of Pharmacy and Medical Sciences, Division of Health Sciences, University of South Australia, SA, Australia; ACRF Cancer Genomics Facility, Centre for Cancer Biology, SA Pathology, Frome Road, Adelaide, SA 5000, Australia; CAS-MPG Partner Institute for Computational Biology, Shanghai Institutes for Biological Sciences, Chinese Academy of Sciences, 320 Yueyang Road, Shanghai 200031, China; School of Molecular and Biomedical Science, the University of Adelaide, Adelaide, SA 5000, Australia; Department of Clinical Science, University of Bergen, 5021 Bergen, Norway; Department of Biomedical Informatics (DBMI), Vanderbilt University Medical Center (VUMC), 2525 West End Ave, Suite 800, Nashville, TN 37203, USA; Institute of Immunology, Second Military Medical University, 800 Xiangyin Road, Shanghai 200433, China

## Abstract

Biological and biomedical research relies on comprehensive understanding of protein-coding transcripts. However, the total number of human proteins is still unknown due to the prevalence of alternative splicing and is much larger than the number of human genes.

In this paper, we detected 31,566 novel transcripts with coding potential by filtering our *ab initio* predictions with 50 RNA-seq datasets from diverse tissues/cell lines. PCR followed by MiSeq sequencing showed that at least 84.1% of these predicted novel splice sites could be validated. In contrast to known transcripts, the expression of these novel transcripts were highly tissue-specific. Based on these novel transcripts, at least 36 novel proteins were detected from shotgun proteomics data of 41 breast samples. We also showed L1 retrotransposons have a more significant impact on the origin of new transcripts/genes than previously thought. Furthermore, we found that alternative splicing is extraordinarily widespread for genes involved in specific biological functions like protein binding, nucleoside binding, neuron projection, membrane organization and cell adhesion. In the end, the total number of human transcripts with protein-coding potential was estimated to be at least 204,950.

**Author summary:** The identification of all human proteins is an important and open problem. In this report we first develop an *ab initio* predictor to collect candidate gene models as many as possible. Next, comprehensive sets of RNA-seq data from diverse tissues and cell lines are used to select confident transcripts. Experimental validation of a subset of predictions confirms a high accuracy for the predicted coding transcript set and has added about 30,000 new protein-coding transcripts to the existing corpus of knowledge in this area. This is significant progress given that the existing protein-coding transcript number in public databases is about 60,000. Our newly found transcripts are more tissue specific. Based on our results, we show that L1's high impact on gene origin and genes with high number of transcripts are enriched in specific functions. At last, we estimate that the total number of human protein-coding transcripts is in excess of 200,000.

## Introduction

Comprehensive gene/transcript annotations are critical reference data for biological studies, especially for genome-wide analyses based on genome annotation. However, alternative splicing (AS) increases the diversity of the transcriptome and proteome tremendously[1] and makes the task of creating a comprehensive gene annotation much harder.

AS occurs in organisms from bacteria, archaea to eukarya[2]. Only a few examples can be found in bacteria[3] and archaea[4,5], but AS is ubiquitous in eukarya[2]. Especially, AS is observed at a higher frequency in vertebrate genomes than in invertebrate, plant and fungal genomes[6,7]. In the human genome, the estimated proportion of genes that undergo alternative splicing has been expanded greatly since the start of this century from 38%[8] to 92%-94%[9-11]. The number of human transcripts generated by AS is estimated to reach 150,000 based on mRNA/ESTs[12], which is still underestimated based on the recent data of GENCODE project[13]. Another research based on RNA-seq data shows that there are ~100,000 intermediate- to high-abundance AS events in major human tissues[9]. The GENCODE Project[13] aims to annotate all evidence-based gene features including protein-coding genes, noncoding RNA loci and pseudogenes for human. The GENCODE V19 contains 196,520 transcripts, of which 81,814 are protein-coding transcripts. However, only 57,005 of them are full length transcripts. Two recent large scale human proteome studies[14,15] expand our understanding on this field. With proteomics data from 17 adult tissues, 7 fetal tissues and 6 purified primary haematopoietic cells, a number of novel proteins were newly identified[14]. In our opinion, a very large proportion of alternative isoforms are still missing, considering the low level of MS/MS spectra of human proteins matching proteins in the Refseq[14]. Overall, finding the total number of all transcripts or protein-coding transcripts encoded in the human genome is still an open problem.

RNA-seq technology is a powerful tool to study the transcriptome and many methods have been developed to reconstruct transcripts from RNA-seq data with[16–19] or without[18–24] transcript annotations. Some of these methods[16,18,19] are based on spliced alignment tools[25–30]. The recent RNA-seq Genome Annotation Assessment Project (RGASP)[31,32] has evaluated 25 protocol variants of 14 independent computational methods for exon identification and transcript reconstruction. Most of these methods are able to identify exons with high success rates, but the assembly of full length transcripts is still a great challenge, especially for the complex human transcriptome[31]. Among those protein-coding region(CDS) reconstruction methods, the transcript-level sensitivity of CDS reconstruction is no more than 20%[31],underscoring the difficulty of transcript detection. Methods assembling transcripts from mRNA-seq reads directly are not that reliable[31] and their limitations have been reviewed by Martin[33].

In this paper, we first introduce ALTSCAN (ALTernative splicing SCANner), which is developed to construct a comprehensive protein-coding transcript dataset using genomic sequences only. For each gene locus, it can predict multiple transcripts. We apply it in candidate gene regions in the human genome and 50 RNA-seq datasets from public databases are used to validate the predicted transcripts. Novel validated transcripts are reported and their characteristics are analyzed. In addition, PCR experiments followed by high throughput sequencing are conducted to verify the existence and expression patterns of these novel transcripts. Moreover, based on the novel transcripts, shotgun proteomics data from 36 breast cancer samples and 5 normal samples are used to search for novel peptides. We have also evaluated the impact of L1 retrotransposons on the origin of new transcripts/genes. In the end, the total number of human transcripts with coding potential has been estimated.

## Results

### Transcript prediction with ALTSCAN

ALTSCAN was developed (see Methods and Figure S1 for details) and applied to human genome sequences (upper part of Figure 1). As a result, 320,784 transcripts with complete ORFs from 33,945 loci were predicted. Among them, 298,454 transcripts were from 22,606 loci in GENCODE or Refseq gene regions; 8,331 transcripts were from 2,721 loci overlapped with pseudogenes; and almost all remained transcripts located in repeat-rich regions. Notably, 9,682 transcripts from 7,663 loci overlapped more than 50% (of each transcript) with L1 elements.

**Figure 1.**
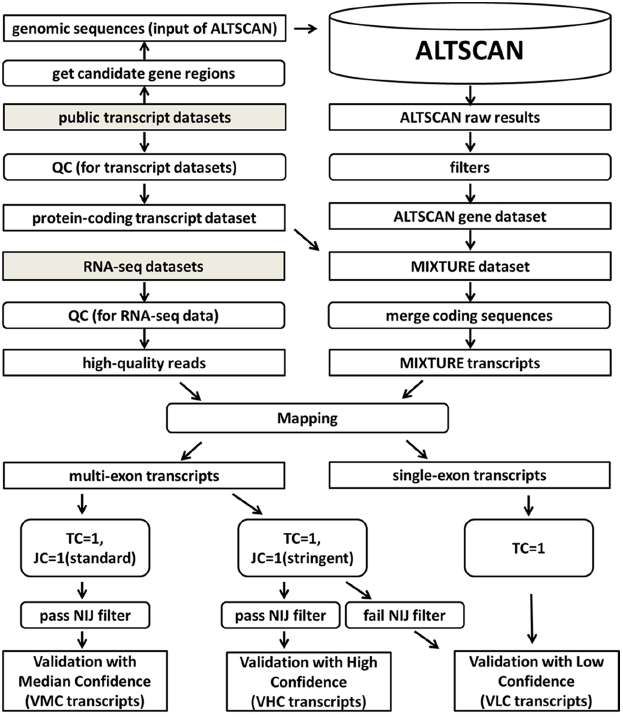
The diagram of transcript prediction using ALTSCAN and validation pipeline based on RNA-seq datasets. The upper part showed the pipeline of alternative transcripts prediction and the MIXTURE dataset construction. The lower part showed the pipeline of transcript validation with RNA-seq data. The grey blocks described raw public data. Candidate gene regions were extracted from various public annotations and then ASs were predicted by ALTSCAN for these regions. Together with the well-annotated KNOWN transcripts, ALTSCAN transcripts were validated with a large number of RNA-seq data. TC was short for transcript coverage and JC was short for junction coverage. The NIJ (novel internal junction) filter was used to check if novel internal junction(s) existed in transcripts (Figure S3). The novel transcript dataset VHC, VMC and VLC were defined as in the figure.

GENCODE and Refseq transcripts were merged to form a dataset named KNOWN (Figure 2). The KNOWN dataset had 2.76 transcripts per gene in average while the number of ALTSCAN dataset was 9.63. 9,780 transcripts from 8,325 genes in ALTSCAN dataset were consistent with the KNOWN dataset and 84.6% of these consistent transcripts were predicted from sub-optimal paths (Figure S2). Next, KNOWN and ALTSCAN dataset were then merged together to form a dataset called MIXTURE. In total, the MIXTURE dataset contained 367,878 transcripts from 28,087 loci. The reduced gene locus number was due to some relatively long transcripts bridging different clusters of transcripts.

**Figure 2.**
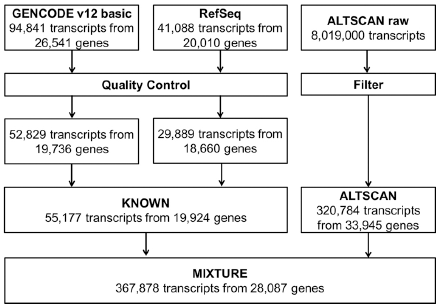
Transcript and gene numbers in dataset construction. Number of transcripts and genes in each dataset was shown. GENCODE and Refseq raw transcripts sharing the same coding regions, having internal stop codons or short introns (<20bp) were removed. Partial-length transcripts were also removed. ALTSCAN raw transcripts sharing the same coding regions were merged and those without complete coding region were filtered out.

Based on the KNOWN dataset, we compared the performance of ALTSCAN with 3 *ab initio* predictors[20,34,35] available in UCSC Genome Browser, as well as 7 predictors[36-39] evaluated in RGASP[31] with capability of predicting coding regions (Table 1). As a result, ALTSCAN’s gene-level sensitivity and specificity were 41.8% and 24.4% respectively, which were much higher than other *ab initio* predictors (the highest one with a sensitivity of 16.8% and a specificity of 14.3%). ALTSCAN’s transcript-level sensitivity and specificity were 17.7% and 3.0% (compared to 6.1% and 14.4% for AUGUSTUS_noRNA, the best *ab initio* predictor in RGASP). This indicated that ALTSCAN could predict many transcripts missed by other *ab initio* predictors. Though the false positive rate of ALTSCAN might be high, we showed that it could be reduced by using RNA-seq data. Integrating RNA-seq data could improve the performance greatly, which could be inferred from the comparison of the performance of AUGUSTUS with and without RNA-seq data. However, ALTSCAN’s gene- and transcript-level sensitivities are even comparable to the best predictor using RNA-seq data. ALTSCAN’s strategy was to filter the predicted transcripts with diverse RNA-seq data to reduce the false discovery rate, which would be further evaluated with real-time PCR.

**Table 1.** Assessment of protein coding region prediction based on the KNOWN dataset.

| Predictor type | Predictor | Gene |  | Transcripts |  | Multiple transcripts per gene |
| --- | --- | --- | --- | --- | --- | --- |
|  |  | sensitivity | specificity | sensitivity | specificity |  |
| *ab initio* | ALTSCAN | 41.8% | 24.4% | 17.7% | 3.0% | yes |
|  | Genscan<sup>b</sup> | 11.3% | 2.2% | 4.1% | 2.2% | no |
|  | Geneid<sup>b</sup> | 13.0% | 8.0% | 4.7% | 8.1% | no |
|  | AUGUSTUS_noRNA<sup>a</sup> | 16.8% | 14.3% | 6.1% | 14.4% | yes |
| *RNA-seq* | AUGUSTUS_RNA<sup>a</sup> | 41.6% | 49.8% | 16.5% | 44.3% | yes |
|  | Exonerate<sup>a</sup> | 36.1% | 23.3% | 15.3% | 27.3% | no |
|  | mGene<sup>a</sup> | 34.6% | 11.2% | 12.6% | 11.3% | yes |
|  | mGene_graph<sup>a</sup> | 31.5% | 47.6% | 12.6% | 29.4% | yes |
|  | mTim<sup>a</sup> | 17.3% | 53.4% | 6.6% | 43.0% | yes |
|  | NextGeneidAS<sup>a</sup> | 24.8% | 28.8% | 9.6% | 30.9% | no |
|  | NextGeneidAS ab-initial<sup>a</sup> | 24.8% | 26.8% | 9.6% | 28.8% | no |
|  | Transomics<sup>a</sup> | 33.4% | 19.9% | 12.2% | 20.2% | no |
|  | Tromer<sup>a</sup> | 3.5% | 1.5% | 1.3% | 0.6% | yes |

<sup>a</sup>Predictions of these methods were derived from RGASP directly.
<sup>b</sup>Predictions of these methods were downloaded from UCSC Genome Browser.

In addition, we compared the correct predictions from ALTSCAN, AUGUSTUS_RNA, Exonerate, mGene and Transomics and found 36% (3,522/9,780) of ALTSCAN’s predictions could NOT be detected by the other 4 methods. The numbers for AUGUSTUS_RNA, Exonerate, mGene and Transomics were 13% (1,261/9,105), 21% (1,792/8,453), 10% (667/6,977) and 8% (569/6,743) respectively. We made similar comparison among *ab initio* predictors. For those correct predictions, 55% (5,410/9,780) of ALTSCAN, 18% (621/3,369) of AUGUSTUS_noRNA, 15% (401/2,631) of Geneid and 6% (127/2,269) of Genscan transcripts could not be detected by the other 3 methods. Therefore, ALTSCAN could detect many transcripts that other methods missed. It is complementary to current methods.

### RNA-seq validation

We used 26 public datasets (50 RNA-seq runs) to validate MIXTURE transcripts, which could be grouped to 3 subgroups based on data sources and read lengths (GROUP I, II and III, Table S1). These transcriptome data were then applied to validate MIXTURE transcripts. We first checked the validation landscape of KNOWN transcripts. Using the standard strategy, we could validate about 10k~20k multi-exon KNOWN transcripts from each RNA-seq dataset (Figure 3A and Table S2); and totally, 40,797 multi-exon KNOWN transcripts (73.94% of all KNOWN transcripts, or 76.91% of KNOWN multi-exon transcripts) were validated, of which, 36,128 transcripts were validated from at least 2 different datasets (Figure 3B and Table S3). Using the stringent strategy, the number of validated transcripts from each dataset were slightly smaller (Figure 3A and Table S2); totally, 35,037 multi-exon KNOWN transcripts (63.50% of all KNOWN transcripts, or 66.05% of multi-exon KNOWN transcripts) were validated, of which, 29,068 transcripts were validated from at least 2 datasets (Figure 3B and Table S3). 5,429 (15.50% of 35,037) transcripts were validated from a specific tissue alone, which implied their tissue-specific expression. Furthermore, 1,992 single-exon transcripts (63.70% of single-exon KNOWN transcripts) were also validated.

**Figure 3.**
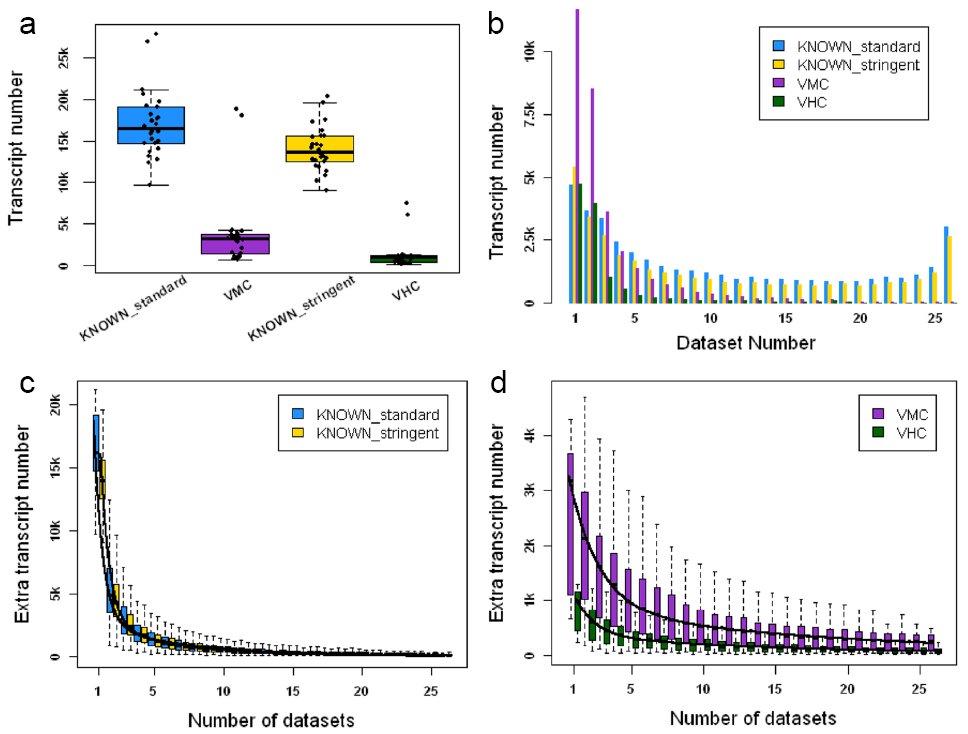
Validation summary of KNOWN and novel transcripts. **A.** showed the number of KNOWN and novel transcripts validated by each RNA-seq dataset using standard or stringent strategy. The highest two points in each group represents validated number from GROUP II datasets (RNA-seq data sequenced from the 16 tissues mixture). **B.** showed validated KNOWN and novel transcript numbers using standard or stringent strategy grouped by numbers of validated datasets. **C** and **D** showed the extra numbers of validated KNOWN and novel transcripts using standard or stringent strategy when a new RNA-seq dataset was added. This process was simulated for 1,000 times with bootstrapping strategy.

Next, we checked the validation landscape of ALTSCAN novel transcripts. Using the standard strategy, 31,819 transcripts were validated with medium confidence (the VMC transcripts). 20,124 of these transcripts were validated from at least 2 datasets. Using the stringent strategy, 11,772 transcripts were validated with high confidence (the VHC transcripts). 7,025 VHC transcripts were validated from at least 2 datasets (Figure 3B and Table S3). 4,747 (40% of 11,772) VHC transcripts were validated from only one dataset. If transcripts validated from less than 5 samples were considered as tissue-specific, we found novel transcripts (VHC or VMC transcripts) had more tissue-specific transcripts than KNOWN (Fisher's exact test, p-values < 0.001). Therefore, the novel transcripts tended to be more tissue-specific (also see Figure 3C and D). In addition, 8,238 transcripts (5,104 single-exon and 3,134 multi-exon transcripts without novel internal junction sites) were also validated as VLC transcripts.

### PCR validation of novel transcripts

We designed primers flanking splice sites of the VMC transcripts, and then randomly selected 88 VMC transcripts (including 32 VHC transcripts) (Table S4). We also designed primers for 8 transcripts of house-keeping genes as positive controls. Real time PCR was applied on 48 samples (tissues or cell lines, Table S5). Then the products from different samples were mixed and sequenced by the Illumina MiSeq platform. As a result, 8 (8/8 = 100%) house-keeping transcripts were validated by at least one sample, indicating the effectiveness of the PCR validation strategy. For the 88 VMC transcripts, 74 were validated by at least one sample, and the success discovery rate achieved 84.1% (74/88 = 84.1%). For the 32 VHC transcripts, 29 were validated by at least one sample, and the success discovery rate was 90.6% (29/32 = 90.6%).

In addition, PCR followed by MiSeq sequencing results showed that the expressions of most of these validated novel transcripts were tissue-specific (Figure 4). For instance, PSMB2 is a gene influences cooperative proteasome assembly [40], homologous recombination [41] and DNA double-strand break repair [41]. Primers were designed to validate the skip of an exon in PSMB2 gene (primer n03 in Figure 4).This exon skipping event was found in 18 tissues and 20 cell lines and the exon was completely skipped in 7 tissues and 13 cell lines (Figure 5). The novel isoform was common in different tissues or cell lines but its expression level was lower than the dominating previous known isoform.

**Figure 4.**
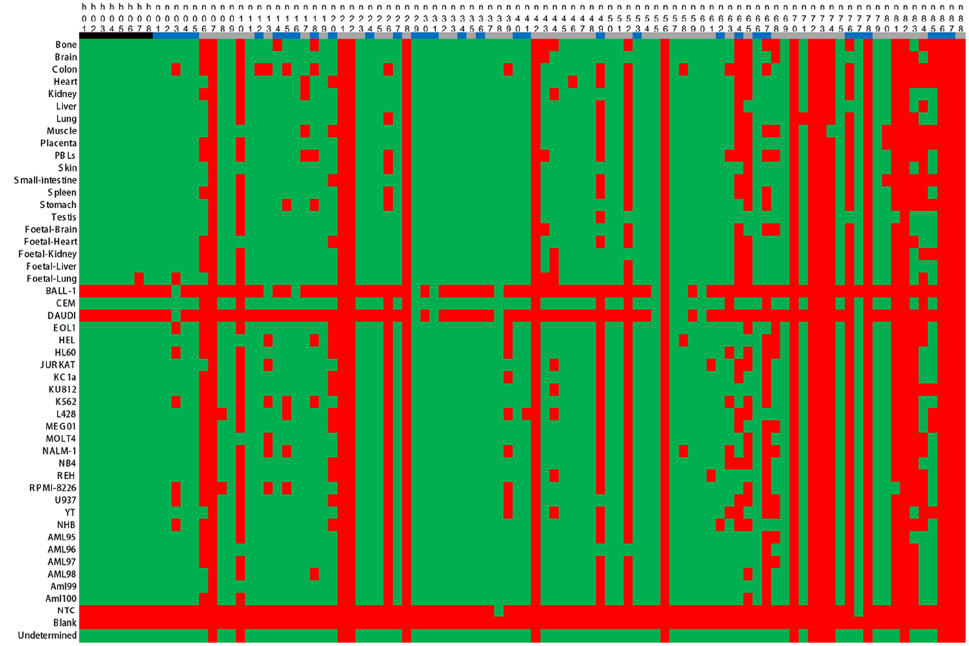
Summary of PCR validation. Black represented house-keeping transcripts; blue represented VHC transcripts; and grey represented transcripts in VMC dataset but not in VHC dataset. Green meant successful validation, while red meant failure. The “blank” line was for a negative control with no RNA used. Reads that failed to be classified clearly by the barcodes were merged to “undetermined”.

**Figure 5.**
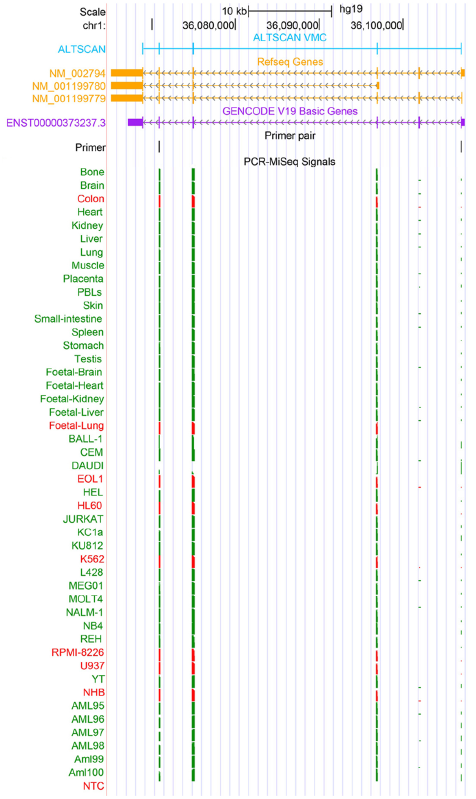
PCR validation of a novel isoform of PSMB2 gene. The exons of ALTSCAN VMC, Refseq and GENCODE transcripts were shown as boxes in light blue, orange and purple respectively. The second coding exon from the 5’ end was skipped in ALTSCAN VMC annotation. Primers used for the validation (forward 5’ CTCCAGACATTTCCTAAGGAGTTC3’ and reverse 5’ CAATATTGTCCAGATGAAGGACGGA3’) were shown in black. MiSeq sequencing results of PCR products were shown as PCR-Miseq signals in green and red. Green indicated the transcript was validated in the tissues or cell line and red meant the transcript was not validated. This novel isoform of PSMB2 gene was validated in most tissues and cell lines except in colon, EOL1, K562and NHB. In HL60, RPMI-8226 and U937 cell lines, it seemed the novel isoform did exist, but the numbers of reads covering the splicing sites were not big enough to meet the validation criteria. In fetal-lung, NM_001199780 from Refseq annotation seemed to be the only expressed isoform.

### Detection of novel proteins

The VHC/VMC transcripts held complete ORFs and therefore had coding potential. Here we used shotgun proteomics datasets from 36 breast cancer samples and 5 normal breast samples to validate the coding potential of these transcripts. The proteomics datasets were used to search against a protein database combining Refseq and the VMC transcripts. Candidate novel peptides from VMC transcripts only were further filtered with GENCODE and Swiss-Prot[42] proteins. As a result, 36 novel proteins supported by at least 2 different peptides including at least 1 novel peptide were detected (Table S8). For instance, we detected two novel peptides encoded in the intron of AEBP2 gene (Figure. 6A). Moreover, 23 of these 36 novel proteins had at least one novel peptide covering novel splice junction sites. For instance, we detected a novel isoform for STUB1 gene (Figure 6B and C). STUB1 protein, a member of E3 ubiquitin ligase, works as a link between the chaperone (heat shock protein 70/90) and proteasome systems[43]. It is also found to be involved in neurodegenerative diseases[44] and cancers[45]. The novel peptide came from the exon-exon junction of the 5^th^ and 6^th^ coding exons, where alternative donor sites were found. As a consequence, 6 amino acids between the tetratricopeptide-like helical domain and the U box domain were removed from the previously known protein. This novel peptide was only detected from cancer samples. It may be a functional isoform related to cancers.

**Figure 6.**
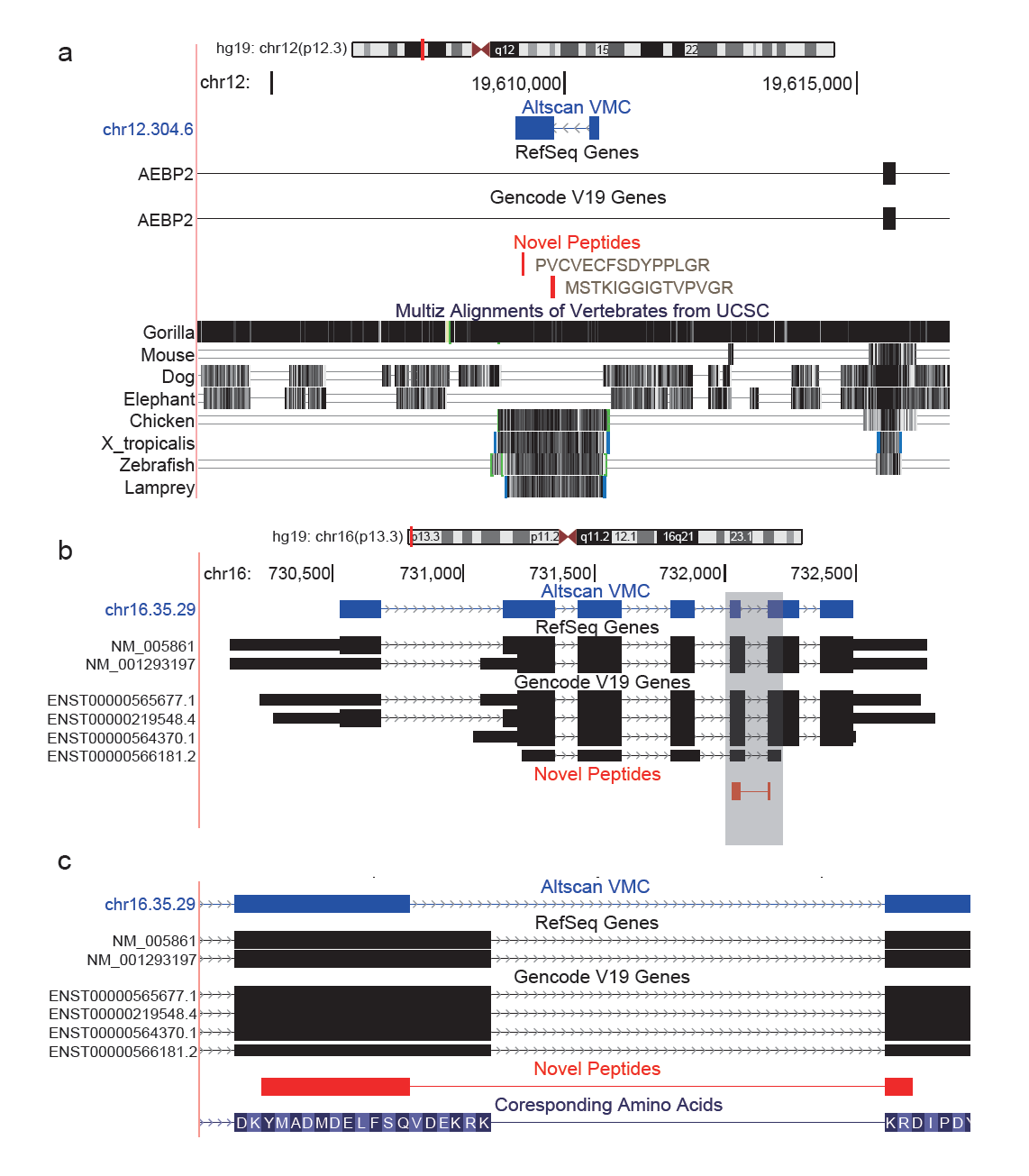
Illustration of novel proteins. **A.** Two novel peptides encoded by a novel gene in the intron of AEBP2 gene. “PVCVECFSDYPPLGR” was detected for 4 times and “MSTKIGGIGTVPVGR” for once. **B.** Novel peptides detected for STUB1 gene. c. Enlarged view of the novel junction (gray area of **B**). The novel peptide “YMADMDELFSQKR” was detected for 5 times. Compared to the GENCODE/Refseq protein, 6 amino acids between the tetratricopeptide-like helical domain and the U box domain were removed.

### Exploring novel genes

Most of the transcripts in VHC or VMC transcripts were novel isoforms of KNOWN genes. However, 1,053 VMC transcripts from 673 loci (including 485 VHC transcripts from 351 loci) were found out of KNOWN gene regions (see Methods and Supplementary material). 782 VMC transcripts from 594 loci (including 312 VHC transcripts from 266 loci) remained after the pseudogenes were removed. Almost all the remained transcripts overlapped with L1 repeat elements. 583 VMC transcripts from 442 loci (including 257 VHC transcripts from 224 loci) were fully covered by single L1 elements (Figure S5A-B). It was reported that a small number of human-specific L1 elements remained retrotransposition-competent[46] and undergo AS[47], and these novel transcripts might be the product of active L1 repeat elements. In addition, 154 VMC transcripts from 128 loci (including 40 VHC transcripts from 32 loci) overlapped partially with L1 elements. 10 out of the 40 VHC transcripts extended out of L1 regions (Figure S5C), indicating their capacity of attacking other genes. The remained 30 VHC transcripts bridged two or more repeat elements, including LINEs, SINEs and LTRs (Figure S5D). These repeat elements expanded the complexity of splicing, which was also known as exonization[48]. Transcripts overlapping partially with L1 elements were at the very early stage towards the well-defined functional transcripts and might be likely dropped in the process of evolution[49]. We provided hundreds of such “young” transcripts. The remained 15 VHC transcripts from 10 loci didn’t overlap with L1 elements (Figure S6). 6 out of the 10 genes shared the same splice sites annotated as non-coding RNAs previously. However, we found complete ORFs in them, suggesting their coding potential. Recent human proteome studies also showed direct evidence that non-coding RNAs can encode peptides[14,15]. One of the 10 genes were absent from GENCODE V12 annotation but were added in the V17 version, while the splicing pattern we provided was different. Another one of the 10 genes was conserved among primates and some non-placental vertebrates in its coding region. The remaining two genes located in the intron or UTR region of known genes. Similar novel coding regions were also found in recent human proteome studies[14].

### AS events analysis

Recent RNA-seq analysis indicated that 95% of human multi-exon genes are alternatively spliced[11]. However, up to now, there are still 5,166 multi-exon genes with only one transcript in KNOWN dataset. We introduced 31,566 VMC/11,549 VHC transcripts (pseudo-transcript removed), which increased the average transcript number per gene from 2.76 to 4.18/3.30 and decreased the proportion of multi-exon genes with single transcript from 30.5% to 25.6%/27.2% (Figure 7A).

**Figure 7.**
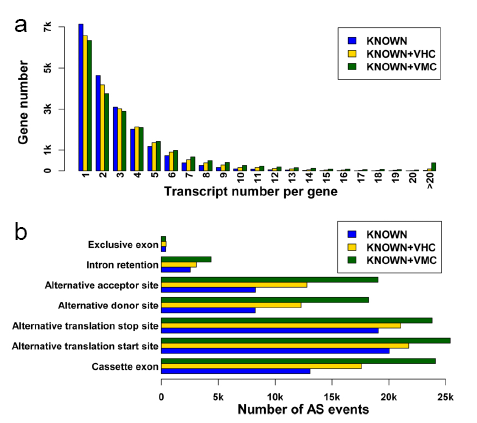
Distribution of alternative splicing in KNOWN and novel datasets. **A.** Distributions of transcript number per gene in KNOWN, KNOWN+VHC and KNOWN+VMC datasets. Genes are grouped by their transcript number. X-axis stands for the group (the number of transcripts per gene), and Y-axis stands for the numbers of genes in each group. All genes with transcript numbers more than 20 were merged in the same group. **B.** The number of different AS events in KNOWN, KNOWN + VHC and KNOWN+VMC datasets. In order to be comparable, number of AS events involved in each type were measured by number of splice sites (see Supplementary material for details).

We checked the splicing patterns for the validated transcripts. Since our research focused on coding regions, those AS events out of coding regions were ignored. Among all splicing patterns, alternative translation start site contributed the most to the complexity of human proteome as described in KNOWN, KNOWN+VHC and KNOWN+VMC datasets (Figure 6B and Table S6). However, alternative translation start sites and alternative translation stop sites, similar with alternative promoter and alternative polyA, are mainly induced by transcription regulation instead of splicing regulation[50]. Ignoring alternative translation start or stop sites, exon skipping accounts for most, which is consistent with our knowledge[11,50]. Compared with KNOWN transcripts, we found that exon skipping, alternative donor sites and alternative acceptor sites accounted for even more proportion in KNOWN+VMC or KNOWN+VHC transcripts (p-values of Fisher’s exact test <0.001). Alternative splice acceptor or donor sites were known to be an intermediate state between constitutive and alternative cassette exons, therefore might be prevalent in human proteome[7].

### Functional analysis

GO (Gene Ontology) enrichment analysis is widely used in biological studies and the background distribution of GO functions is critical in analysis procedures. We carried out GO annotation for these novel transcripts. As a result, the function distribution of the VHC/VMC transcripts was quite consistent with that of the KNOWN transcripts (Figure S8A-F). The Pearson correlation coefficients of function distribution between VHC and KNOWN transcripts were 0.985, 0.950 and 0.967 in biological process, molecular function and cellular component level respectively (Figure S8G-I). The corresponding coefficients between VMC and KNOWN transcripts were 0.989, 0.970 and 0.988, respectively (Figure S8J-K). These results indicated that novel transcripts predicted by our methods had similar function distribution with known transcripts.

**Figure 8.**
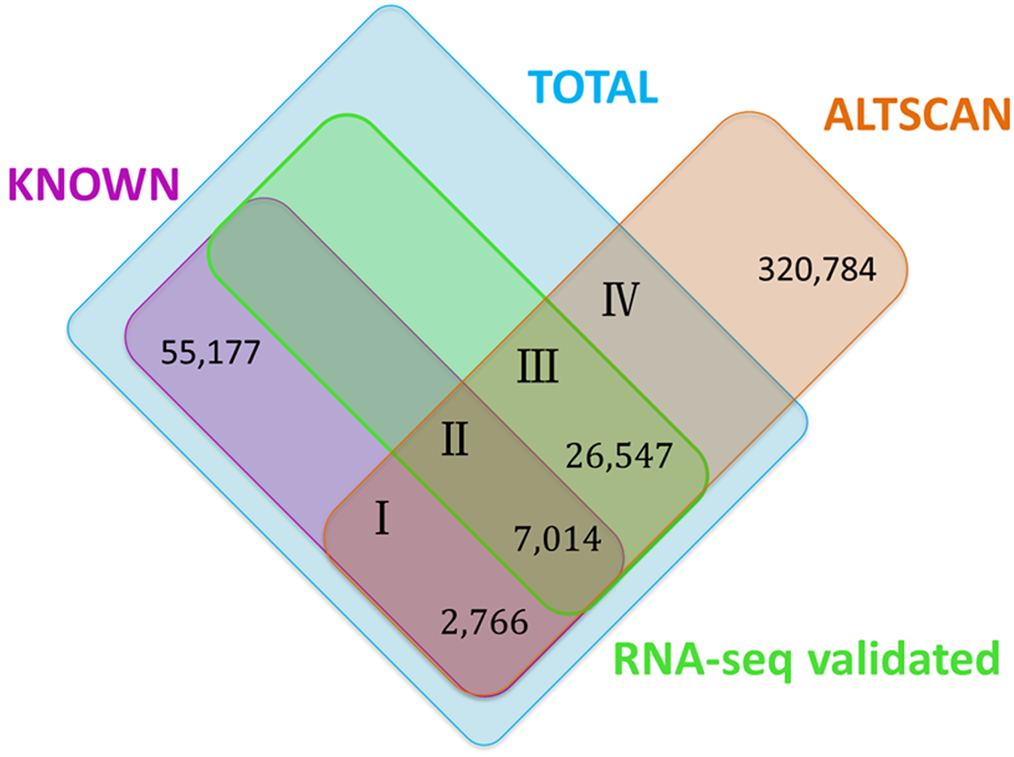
Estimation of the number of transcripts with coding potential. Datasets are illustrated in different colors. “All” means the total transcript dataset whose transcript number was to be estimated. Different datasets were represented by different numbers. *I* represents transcripts in KNOWN and ALTSCAN datasets but haven’t been validated by RNA-seq data used in this study; *II* represents transcripts in KNOWN and ALTSCAN datasets and have been validated by RNA-seq data used in this study; *III* represents VHC or VHC+VMC transcripts; *IV*represents novel but real transcripts in ALTSCAN datasets that have not been validated by RNA-seq data used in this study.

In order to investigate influence of AS on specific biological process, we examined whether transcript numbers of genes were related with biological functions and pathways. Enrichment analysis of genes with high number (>5) of transcripts showed AS is enriched in specific molecular functions (binding, especially protein binding and nucleoside binding and enzyme regulatory activities like transcription cofactor activity), cellular components (neuron, membrane-related locations and cell junction) and biological process (regulation of small GTPase mediated signal transduction, vesicle-mediated transport and membrane invagination) (Table S7). However, no enriched KEGG pathways was found. The reason might be that although one gene with different biological functions had AS bias, the overall AS bias for all genes involved in the pathways was not significant.

### Estimation of the total number of transcripts with coding potential

In spite of the advancement of RNA-seq technology, estimating the total number of protein-coding transcripts in human is still an open problem. Many transcripts are expressed at low levels or in a temporally and spatially specific way. As a consequence, they are difficult to be discovered and it is difficult to estimate the total number of human proteins as well. ALTSCAN can be used as an *ab initio* predictor and its sensitivity is irrelevant to expression levels. Therefore, we may assume that ALTSCAN’s sensitivity calculated based on known transcripts is equal to the value calculated by considering the undiscovered transcripts (see Methods for details). Based on this assumption, we estimated that the number of human transcripts with coding potential to be at least 204,950.

## Discussion

AS expands the functional repertoire of human genome, but only a small proportion of AS has been experimentally characterized. A comprehensive gene annotation is critical for genome-wide analysis, cis-regulatory element finding, hereditary disease studies and nearly all biological science studies. Detecting all genes and transcripts for human and other model organisms is one of long term goals of biological research, which can help reveal the essence of life. In this paper, we have introduced ALTSCAN, and demonstrated that by predicting many transcripts in a single locus from the genomic sequence directly and filter the predictions with RNA-seq data, we can generate a big number of novel protein-coding transcripts.

ALTSCAN’s transcript-level sensitivity is 17.7% while the corresponding number is 6.1% for the best existing *ab initio* predictor. It demonstrates that ALTSCAN’s multi-layer Viterbi algorithm is able to detect more transcripts. Recent RGASP assessed many transcript reconstruction methods[31] and the predictions from different methods have been evaluated with expressed transcripts from GENCODE v3c only, instead of all known transcripts in public databases. In our evaluation, the KNOWN dataset (GENCODE and Refseq dataset) was used as the annotation dataset. In RGASP, among all tools, the best transcript level sensitivity for CDS reconstruction was 19.8% (16.5% in our evaluation results, due to the increased number of annotated transcripts by merging the Refseq and GENCODE datasets). This shows that our comparison is reasonable. By comparing the correct predictions from different programs, we have found that ALTSCAN can detect many transcripts that other methods may miss. Therefore, ALTSCAN is complementary to existing methods. Recently, single molecule real-time (SMRT) sequencing was utilized to obtain transcriptome data of 20 human organs and tissues[51,52]. From these transcriptome data, 11,833 transcripts not included in GENCODE were created (from authors Tilgner H. and Snyder MP. [51,52]). 11,084 of them were labeled as “protein-coding”. We compared the VMC transcripts with these 11,084 novel protein-coding transcripts. As a result, 2,214 VMC transcripts were supported, which meant all the splice junctions of a VMC transcript were consistent with a SMRT transcript. The sensitivity on this SMRT novel transcript data was about 20% (2,214/11,084), which was similar to ALTSCAN’s performance in KNOWN dataset. The conservation level of transcripts in VMC, Refseq, GENCODE and novel SMRT transcripts were similar when they were compared to the mouse genome (mm10). It indicated that our assumption to estimate the overall number of human transcripts was somewhat reasonable.

Despite its surprising capability of detecting novel transcripts with high confidence when integrated with RNA-seq data, some limitations existed. First, our results from ALTSCAN were still far from “exhaustion” due to the limitation of algorithm and computing capability. In our extended Viterbi algorithm, the average transcript number discovered had no sign of going down even at a depth of 250, which suggested that this depth was still not enough. Moreover, the initial ALTSCAN prediction before the RNA-seq filter contained many redundant transcripts.

In addition, our RNA-seq studies focused on validation of candidate transcripts without exploring the whole expression profiles in different tissues. Relative strict criteria were used to remove the mapping errors of RNA-seq reads to the reference genome. Recent data of the ENCODE project indicated that about three-quarters of the human genome was capable of being transcribed[53], which increased the importance of mapping of splice junction reads when validating spliced gene structures. Therefore, we paid more attention to the validation of junction sites instead of the “transcription”. In order to get reliable prediction results, the sequencing depth and different parameters in our validation pipeline are assessed for their impact on the number of validated transcripts. Results showed that shorter reads required more strict validation parameters, and deeper sequencing depth could help validate more novel transcripts.

We also found that hundreds of transcribed L1 elements may be still active. L1 elements provided many potential splice sites[54]. After their insertion to new locations of the genome, they could alter the coding potential of nearby nucleotides with their active splice sites. Although it might break a nearby gene, it was a tremendous source of exonization and a driving power of evolution. In addition, we detected several novel proteins encoded by L1 elements in both cancer and normal samples (Table S8).

The identification of all human proteins is an important and unsolved question. Our novel transcripts can help detect novel proteins. Mass spectrometry (MS) and ribosome profiling (RP)[55] method can be utilized to study the proteome. MS method detected peptide segments from a candidate protein pool; and RP method provided only short portions of RNAs that were bound to ribosomes. Recent human proteome studies took a big step towards annotation of all human proteins, however, it was far away from complete, mostly due to isoforms derived from AS, which often differed only several peptides nearby the corresponding splice sites. Therefore they were very difficult to be discovered by both methods[49]. We have detected 62 novel proteins missing in Refseq. 29 of these 62 proteins have novel peptides covering splice junctions. Overall, 9 of the 62 proteins have been annotated in both GENCODE and Swiss-Prot. Among the 62 proteins, 5 and 11 of them have been annotated in GENCODE only and Swiss-Prot only respectively. Therefore, the final number of novel proteins is 61-9-5-11=36. To our knowledge, finding 36 novel proteins in one tissue (41 samples) is quite effective. Surprisingly, 24 of the novel proteins have novel peptides covering novel splice junctions, indicating the capability of our method to detect novel transcripts especially for those with novel splice sites. In short, our work is an effective supplement to existing methods and will help to build a more comprehensive human protein-coding gene annotation.

To conclude, we have developed a novel system to predict protein-coding transcripts by integrating *ab initio* prediction and filtering with RNA-seq data; andwe have detected and validated 11,549~31,566 transcripts with complete ORFs at a FDR of 9.38%~15.9%. In contrast to known transcripts, these novel transcripts are highly tissue-specific. We estimate the total number of full length transcripts to be no less than 200 thousand, which indicated that majority of the protein-coding transcripts are still missing in the current databases. In addition, 36 novel proteins are detected. Furthermore, we find that L1 elements have a far greater impact on the origin of new transcripts/genes than previously thought. Alternative splicing is extraordinarily widespread for genes involved in some basic biological functions.

## Materials and methods

Detailed methods can be found in Supplemental material. Here we described materials and methods briefly.

### ALTSCAN

ALTSCAN utilized an extended Viterbi algorithm. The top N value(s) were kept in each step so that the top N path(s) would be generated, which enabled the scanner to predict multiple transcripts for one gene. N was set to 250 for most ALTSCAN inputs. Figure S1 showed how extended Viterbi algorithm worked.

### ALTSCAN prediction for the human genome

In practice, candidate gene regions were extracted from the human genome as the input to ALTSCAN (upper part of Figure 1). The candidate gene region included the regions of known genes, SIB genes, and NSCAN predicted genes. The known genes included GENCODE basic V12 genes, which were derived from HAVANA manual annotation process and Ensembl automatic annotation pipeline and Refseq genes[56]. SIB genes[57] were genes with support evidences of at least one GenBank full length RNA sequence, one Refseq RNA, or one spliced EST. SIB genes were used to create regions with mRNA or EST evidences. In addition, NSCAN predicted genes were those predicted genes with multiple-genomes. GTF files were collected for all these gene datasets, and a totally, 33,480 sequences including a padding length of 5,000 bts both downstream and upstream of genes were extracted from human genome (hg19). ALTSCAN was run on these regions and raw results were filtered and clustered to ensure each transcript had a unique coding sequence. Finally, 320,784 transcripts with unique complete coding regions from 33,945 genes made up the ALTSCAN prediction for the human genome. Details of ALTSCAN’s prediction on human genome were described in Supplemental material.

### Assessment of coding region (CDS) prediction

We evaluated the performance of tools for CDS prediction including 4 *ab initio* predictors (ALTSCAN, Genscan[35], Geneid[34] and AUGUSTUS[20]) and 7 predictors using RNA-seq data (AUGUSTUS[37], Exonerate[38], mGene[36], mTim, NextGeneid, Transomics and Tromer[39]) based on the KNOWN annotation. Predictions from AUGUSTUS_no_RNA and all predictors using RNA-seq data were downloaded from RGASP[31,32]. The evaluation on gene-, transcript- and exon-level was achieved with the tool RGASP.jar provided by RGASP.

### RNA-seq validation

We collected 50 RNA-seq runs from the Illumina Human BodyMap2 project and ENCODE project. Different runs of a biological sample were merged to 26 datasets. These datasets were further classified into to 3 groups based on data source and sequencing features (see Table S1). We created a pipeline (lower part of Figure 1) to validate known and predicted transcripts with these RNA-seq data. Quality control of RNA-seq data were processed using the NGSQC[58]. Coding sequences from MIXTURE transcripts were extracted with 100nts upstream start codons and 100nts downstream stop codons. These coding fragments formed the mature transcript dataset. High quality reads were mapped to mature transcript dataset using Bowtie[59]. A splicing junction site was covered if and only if at least *M* read(s) covered both sides of the adjacent exons with no less than *L* nts on each side. We used two strategies in our splice junction site validation: the standard strategy (*L*=10 and *M*=1) and the stringent strategy (*M*>5 and *L*>7, Figure S4). In addition, novel validated transcripts (in ALTSCAN but not in KNOWN dataset) were further filtered by the NIJ (novel internal splice junction, Figure S4) filter and grouped into VHC (validation with high confidence), VMC (validation with median confidence) and VLC (validation with low confidence) datasets (Figure 1).

### PCR validation of novel transcripts

Primers were designed with Primer3[60]. Real-time PCR was conducted using EvaGreen on the Biomark System (Fluidigm). PCR products from the same samples were mixed and barcodes were added. Finally, samples were pooled and sequenced using Illumina MiSeq sequencer (see PCR experiment part in Supplementary material).

### Detection of novel proteins

Shotgun proteomics data of 36 breast cancer samples (900 raw files) and 5 normal breast samples (125 raw files) downloaded from CPTAC were used in this study[61]. The mass spectrometry raw data were searched against a combined database including Refseq protein sequences, VMC protein sequences and a decoy database with all protein sequences reversed, using the X!Tandem search engine[62]. The false discovery rate (FDR) was set at 10^-6^ as previously described[63]. Peptides that could be scored according to the VMC transcripts but could not be scored according to the Refseq transcripts were identified as the preliminary novel peptides. Proteins that could be mapped by at least two identified unique peptides including at least one novel peptide were defined as candidate novel proteins. These preliminary peptides were further aligned to GENCODE (version 12) and Swiss-Prot[42] (downloaded on Dec. 1, 2014) proteins to get the final novel peptides using NCBI BLAST (blastp).

### AS event analysis

AS events were classified into seven categories and were detected with methods described in Supplemental material and Figure S7.

### Functional analysis

Enrichment analysis was carried out with DAVID[64] and iGepros website[65]. Enrichment p-values were adjusted with Benjamini-Hochberg method.

### Estimation of the total number of transcripts with coding potential in human

In order to estimate the total number of transcripts with coding potential in human, we assumed the sensitivity of ALTSCAN evaluated by known transcript was equal to the value calculated with the consideration of undiscovered transcript. The relationship between the datasets (I, II, III and IV) is shown in Figure 8.ALTSCAN’s sensitivity evaluated by known transcript was calculated as

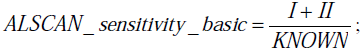

ALTSCAN’s sensitivity considering undiscovered transcripts can be described as

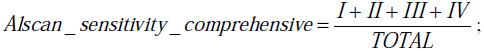

then the total number of transcripts with coding potential in human can be described as

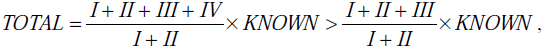

where *I* + *II* = 9,780, and *III* = 31,566 × 84.1% = 26,547 (VMC transcript number multiplied by accuracy estimated from PCR validation). represents novel transcripts predicted by ALTSCAN without RNA-seq *IV* validation. We found that using GROUP II data only (sequenced from mixtures of 16 tissues), 30,433 VMC transcripts could be obtained. The other 24 datasets contributed extra 1,133 transcripts; It indicated *IV* would be a small proportion of the total transcripts.

## Acknowledgements

We thank the High Performance Computing Center (HPCC) at Shanghai Jiao Tong University for the computation. We thank Dr. Hagen Tilgner and Dr. Michael Snyder for providing transcript data with PacBio sequencing support. We thank Dr. Guohui Ding from Chinese Academy of Science and Dr. Yuanyuan Li from Shanghai Center for Bioinformation Technology for their helpful discussion and insightful comments.

## Supporting Information

**Supplemental material**. Supplemental information including supplemental methods, figures and tables.

